# MAVS regulates the quality of the antibody response to West-Nile Virus

**DOI:** 10.1101/2019.12.15.875906

**Authors:** Marvin O’Ketch, Cameron Larson, Spencer Williams, Jennifer L. Uhrlaub, Rachel Wong, Neha R. Deshpande, Dominik Schenten

**Affiliations:** Department of Immunobiology, University of Arizona, Tucson, AZ 85724; Division of Biological and Biomedical Sciences, Washington University in St. Louis, Saint Louis, MO 63110

## Abstract

A key difference that distinguishes viral infections from protein immunizations is the recognition of viral nucleic acids by cytosolic pattern recognition receptors (PRRs) such as RNA-sensing Rig-I-like receptors (RLRs). Insights into the specific functions of cytosolic PRRs in the instruction of adaptive immunity are therefore critical for the understanding of protective immunity to infections. West Nile virus (WNV) infection of mice deficent of MAVS, the essential RLR signaling adaptor, results in a defective adaptive immune response. While this finding suggests a role for RLRs in the instruction of adaptive immunity to WNV, it is difficult to interpret due to the high WNV viremia, associated exessive antigen loads, and pathology in the absence of a MAVS-dependent innate immune response. To overcome these limitations, we have infected MAVS-deficient mice with a single-round-of-infection mutant of WNV called RepliVAX (RWN). RWN-infected MAVS-deficient (MAVS^KO^) mice failed to produce an effective neutralizing antibody response to WNV despite normal titers of antibodies targeting the viral WNV-E protein. This defect occurred indepedently of antigen loads or overt pathology. The specificity of the antibody response in RWN-infected MAVS^KO^ mice remained unchanged and was still dominated by antibodies that bound the neutralizing lateral ridge (LR) epitope in the DIII domain of WNV-E. Instead, MAVS^KO^ mice produced IgM antibodies, the dominant isotype controlling primary WNV infection, with lower affinity for the DIII domain. Our findings suggest that RLR-dependent signals are important for the quality of the humoral immune response to WNV.

## Introduction

The features that separate protective adaptive immune responses from similar responses that fail to protect from infection have yet to be clearly delineated. Aside from important factors such as antigen structure or antibody specificity, it is widely believed that many aspects that define protective immunity are instructed by the innate immune system (1–4). Presumably, pathogens and live vaccines trigger multiple pattern recognition receptors (PRRs) that induce the optimal set of signaling molecules for the regulation of protective adaptive immune responses. In contrast, adjuvant-based subunit vaccines likely represent incomplete mimics of live vaccines that fail to replicate the necessary set of regulatory signals. As the nature and functions of these signals are incompletely understood, rational vaccine design is still facing considerable challenges. Consistent with this view, the highly successful live yellow fever vaccine YF-17D activates multiple PRRs that collectively define the cytokine profile and magnitude of both CD4 and CD8 T cell responses as well as antibody responses (5–7). Likewise, recognition of RNAs uniquely associated with live bacteria can promote the magnitude and vaccine efficacy of T-dependent antibody responses (8–10). Vaccination with live attenuated microbes is therefore still often considered the best way to elicit effective long-lasting cellular and humoral immunity (4, 11–13).

A key feature that distinguishes viral infections from immunizations with subunit vaccines is the activation of cytosolic RNA or DNA-sensing PRRs during the course of viral infections. The Rig-I-like receptor (RLR) family of PRRs include the ubiquitously-expressed RNA-sensing helicases RIG-I and MDA5, which recognize microbial RNA in the cytosol (14). Both receptors rely on the adaptor protein MAVS for the transmission of their signal and the induction of proinflammatory cytokines and interferon responses (14–16). RLRs clearly play an essential function in the regulation of innate immunity to many RNA viruses as mice deficient of components of the RLR signaling pathway often suffer from uncontrolled viral replication and succumb to the infection. However, a clear understanding of RLR function in adaptive immunity has remained elusive as the viral uncontrolled replication complicates efforts to separate intrinsic functions of RLRs from the confounding variables of antigen load and pathology.

Infection of MAVS-deficient mice (MAVS^KO^ mice) with pathogenic West-Nile Virus (WNV), a single-stranded RNA virus of the flavivirus family, results a dysregulated adaptive immune response (17, 18). Specifically, MAVS^KO^ mice generate poorly neutralizing antibodies against WNV, even though they have higher WNV-specific antibody titers than wild-type controls, suggesting that MAVS may play a role in quality control of the antibody response to WNV (17). This finding was surprising because WNV-mediated TLR activation should be sufficient for the generation of humoral immunity to WNV. However, interpreting this result as evidence for a direct link between MAVS-induced signals and the quality of neutralizing antibody responses is challenging because MAVS deficiency also leads to a significant increase in WNV viremia (>1000-fold) that causes severe pathology and death as well as an excess of viral antigens (17). It remains therefore unclear whether MAVS directly contributes to humoral immunity to WNV.

To overcome the limitations of high viral titers associated with WNV infections of mice with deficiencies in innate signaling pathways, we infected MAVS^KO^ mice with a single-round-of-infection mutant of WNV called WNV-RepliVAX (RWN) (19). This mutant virus carries a deletion in the capsid gene and fails to generate infectious viral particles but produces all other viral proteins and RNA. Using this system, we show here that MAVS^KO^ mice fail to generate effective neutralizing antibody responses to RWN even under conditions of similar antigen loads. We show that this defect is caused a T-dependent antibody response that is directed at the neutralizing epitope of WNV but displays a lower affinity for this epitope. Our data therefore suggest that MAVS-dependent signals directly influence the quality of the antiviral antibody response against WNV by calibrating the affinity of the neutralizing antibodies.

## Material and Methods

### Mice

MAVS^KO^ mice, MHCII^KO^ mice, and Rag2^KO^ mice were kept on a C57BL/6 background under SPF conditions (20–22). Experimental mice and wild-type controls were cohoused immediately following weaning. The mice were analyzed between 7-12 weeks of age and involved both sexes. All experiments were performed in accordance with the Institutional Animal Care and Use Committee (IACUC) of the University of Arizona.

### Virus production and mouse infections or immunizations

The single-round-of-infection mutant of WNV (WNN-RepliVAX, RWN) has been described before (23). This WNV-NY-derived virus carries an inactivated capsid gene and requires passaging over capsid-expressing BHK cells for the production of infectious virions. BHK cells were infected with 0.05 MOI of RWN and the supernatants were harvested 48-96 hrs later. RWN titers were determined by infecting fresh Vero cells with serial dilutions of RWN and subsequent intracellular staining for infected cells with a biotinylated humanized anti-WNV-E antibody (hE16-biotin) followed by Streptavidin-horseradish peroxidase (HRP) (24) or, alternatively, with purified hE16, followed by anti-human IgG2a antibody. HRP-positive cells were detected with the TrueBlue substrate. Unless otherwise noted, mice were infected with RWN subcutaneously in the footpads with a dose of 1 x 10^5^ Pfu per mouse. When indicated, mice were infected with 5 x 10^5^ Pfu RWN and 1x 10^6^ Pfu RWN or immunized with 50 μg Ovalbumin and 5 μg LPS in Incomplete Freund’s Adjuvant (all Sigma Aldrich, St. Louis, MO).

### Antibodies and other reagents

Antibodies against CD3ε, CD4, CD8, CD11b, CD11c, CD25, CXCR5, PD-1, CD19, B220, and FoxP3 were purchased from BD Biosciences (San Diego, CA), ThermoFisher (Waltham, MA), or Biolegend (San Diego, CA). PNA was obtained from Vector Laboratories (Burlingame, CA). E641:I-A^b^ tetramers and recombinant DIII and DIII-K307E/T330I were kindly provided by M. Kuhns (University of Arizona) and D. Bhattacharya (University of Arizona), respectively (25, 26). Anti-PE microbeads were purchased from Miltenyi Biotec (Auburn, CA).

### Surface and intracellular staining

Cells were stained with indicated antibodies for 15 min on ice for cell surface staining. For CXCR5 and E641:I-A^b^ tetramer staining, cells were incubated for 45 min at room temperature. E641:I-A^b+^ cells were subsequently enriched with anti-PE-microbeads according to manufacturer’s instructions. Staining for intracellular antigens were performed with the BD Bioscience or Thermofisher (for FoxP3) intracellular staining kits. Cells were analyzed on a LSRFortessa flow cytometer (BD Bioscience, San Diego, CA) and the FlowJo software (Tree Star).

### Detection of WNV-E protein

For the detection of WNV-E antigen levels in RWN-infected mice, 1-2 x 10^7^ cells from the dLNs were isolated 24 hrs after infection and cultured for an additional 24 hrs *in vitro*. Subsequently, supernatants and cells were subjected to one freeze-thaw cycle at −80 °C to release RWN virions from the infected cells. After a centrifugation step, the supernatants were incubated with anti-mouse IgG2a-coupled LEGENDplex beads (Biolegend) that had been coated with the anti-DII/III E60 antibody at a concentration of 1 μg/100 μl of beads. Binding of WNV-E protein to the beads was detected by flow cytometry with the biotinylated anti-DIII hE16 antibody, followed by SA-PE. Samples from naïve mice or standards with and without recombinant WNV-E protein were used as controls.

### Quantitative PCR

RNA was isolated from the dLNs following infection with RWN and converted into cDNA. Quantitative PCR was performed using the PerfeCTa SYBR Green Fastmix (Quantabio, Beverly, MA) on a StepOne Real-time PCR system (Applied biosystems, Foster City, CA). PCR products were amplified with the following primer pairs: Cytokines: IL-1β-F 5’-TGAGCACCTTCTTTTCCTTCA and IL-1β-R 5’-TGTTCATCTCGGAGCCTGTA; IL-6-F 5’-GTTCTCTGGGAAATCGTGGA and IL-6-R 5’-TTTCTGCAAGTGCATCATCG; TNF-α-F 5’-CCCCAAAGGGATGAGAAGTT and TNF-α-R 5’-TGGGCTACAGGCTTGTCACT; IFN-α4-F 5’-AGGACAGGAAGGATTTTGGA and IFN-α4-R 5’-GCTGCTGATGGAGGTCATT; IFN-β-F 5’-CACAGCCCTCTCCATCCACT and IFN-β-R 5’-GCATCTTCTCCGTCATCTCG; IFN-λ2-F 5’-CAGAGCCCAGGTCCCCGA and IFN-λ2-R 5’-CACACTTGAGGTCCCGGGT; IFN-λ3-F 5’-CAGAGCCCAAGCCCCCGA and IFN-λ3-R 5’-CTTGAGGTCCCGGAGGAG; IFIT1-F 5’-GCTGAGATGTCACTTCACATGG and IFIT1-R 5’-CACAGTCCATCTCAGCACACT; IFIT2-F 5’-AGTACAACGAGTAAGGAGTCACT and IFIT2-R 5’-AGGCCAGTATGTTGCACATGG; ISG15-F 5’-GGTGTCCGTGACTAACTCCAT and ISG15-R 5’-TGGAAAGGGTAAGACCGTCCT; and RSAD2-F 5’-TGCTGGCTGAGAATAGCATTAGG and RSAD2-R 5’-GCTGAGTGCTGTTCCCATCT; WNV proteins: E-F 5’-GGCTTCCTTGAACGACCTAA and WNV-E-R 5’-CGTGGCCACTGAAACAAAAG; NS1-F 5’-CAACTCAGAATCGCGCTTGG and NS1-R 5’-CTCTCGAGGATTCCATCGCC; and NS4b-F 5’-AACCCGTCTGTGAAGACAGT and NS4b 5’-ATAAGCACGACAACCAACCC.

### IFN Bioassay

**I**FN activity was measured by a standard assay quantifying the protection of cells from cytopathic effects (CPE) of vesicular stomatitis virus (VSV). L929 cells were plated on 96-well tissue culture plates and incubated for 24hrs at 37 °C. IFN-α2 standard was titrated and serum samples were serially diluted before incubation with VSV on confluent L929 cells for 24 hours. The cells were fixed with paraformaldehyde and stained with a solution containing crystal violet dye. The protection of cells from CPE by each serum sample was scored and compared to IFN-α2 standard samples to establish a concentration value for type I IFN in the serum in Units/ml.

### Immunoglobulin ELISA

Detection of WNV-specific antibodies was based on ELISAs with recombinant WNV-E, DIII, or DIII-KT as antigens. Production of these reagents followed published protocols (27, 28). Briefly, DIII proteins were expressed in BL21(DE3) *E. coli* cells, refolded from inclusion bodies by oxidative refolding, and purified by size exclusion. AviTag-DIII protein was biotinylated and purified again by size exclusion. Serial dilutions of serum from RWN-infected mice were applied to plates coated with recombinant WNV-E, DIII; or DIII-KT proteins. Bound RWN-specific antibodies were detected with biotinylated goat anti-mouse IgM, IgG, or IgG2c (Southern Biotech, Birmingham, AL), followed by streptavidin-conjugated horseradish peroxidase (SA-HRP) and TMB substrate (both BD Bioscience, San Diego, CA). Anti-mouse Ig(H+L) (Southern Biotech, Birmingham, AL) and serial dilutions of mouse IgM and IgG2c (Southern Biotech, Birmingham, AL) were used for standards. High-affinity antibodies were measured similarly using plates coated with recombinant DIII alone or diluted 1:3 with BSA. After the initial binding, low affinity antibodies were washed off by incubating the samples for 15 min in presence of increasing amounts of NaSCN before detection with biotinylated goat anti-mouse IgM or IgG, followed by SA-HRP.

### Virus neutralization

The assay to measure the ability of serum to neutralize WNV has been described before (29). Briefly, 2000 pfu/ml RWN were incubated with serial dilutions of serum from RWN-infected mice for 2.5 hrs at room temperature and subsequently used to infect Vero cells. The number of infected cells was assessed 30-48 hrs later by staining for the expression of WNV-E protein using an anti-WNV-E antibody (E16-biotin). Scored was the lowest dilution factor of serum necessary to achieve a reduction of 90% of infected cells compared to cells infected with WNV without prior serum incubation (PRNT90).

### Statistical Analysis

All experiments were performed independently three or more times. Statistical significance was determined with a Mann-Whitney test using the Prism6 software (GraphPad). Number of asterisks represents the extent of significance with respect to the p value.

## Results

### Impaired neutralizing antibody response to RWN in RWN-infected MAVS^KO^ mice

To dissect the function of MAVS in the regulation of humoral immunity, we infected MAVS^KO^ mice and MAVS^WT^ controls with a replication-incompetent mutant of WNV called WNV-RepliVAX (RWN) in (23). This mutant lacks a functional capsid gene and thus fails to produce infectious progeny but generates otherwise all viral proteins and RNA. RWN-infection of MAVS^KO^ mice in the footpads led to a WNV-E-specific IgM and IgG response similar to wild-type levels on day 8 post-infection (Fig. 1A-B). However, sera from RWN-infected MAVS^KO^ failed neutralize the virus effectively when compared to MAVS^WT^ mice (Fig. 1C). Importantly, the sera from MAVS^KO^ mice also showed a neutralization defect of WNV when the amounts of RWN-specific IgM and IgG Abs were taken into account, as MAVS^KO^ mice exhibited a significantly lower neutralization index than MAVS^WT^ mice (neutralization divided by amount of virus-specific IgM + IgG) (Fig 3D). This finding was also true when the neutralization index was calculated based on the IgM or IgG titers alone (Supplementary Fig. S1A-B).

**Figure 1.**
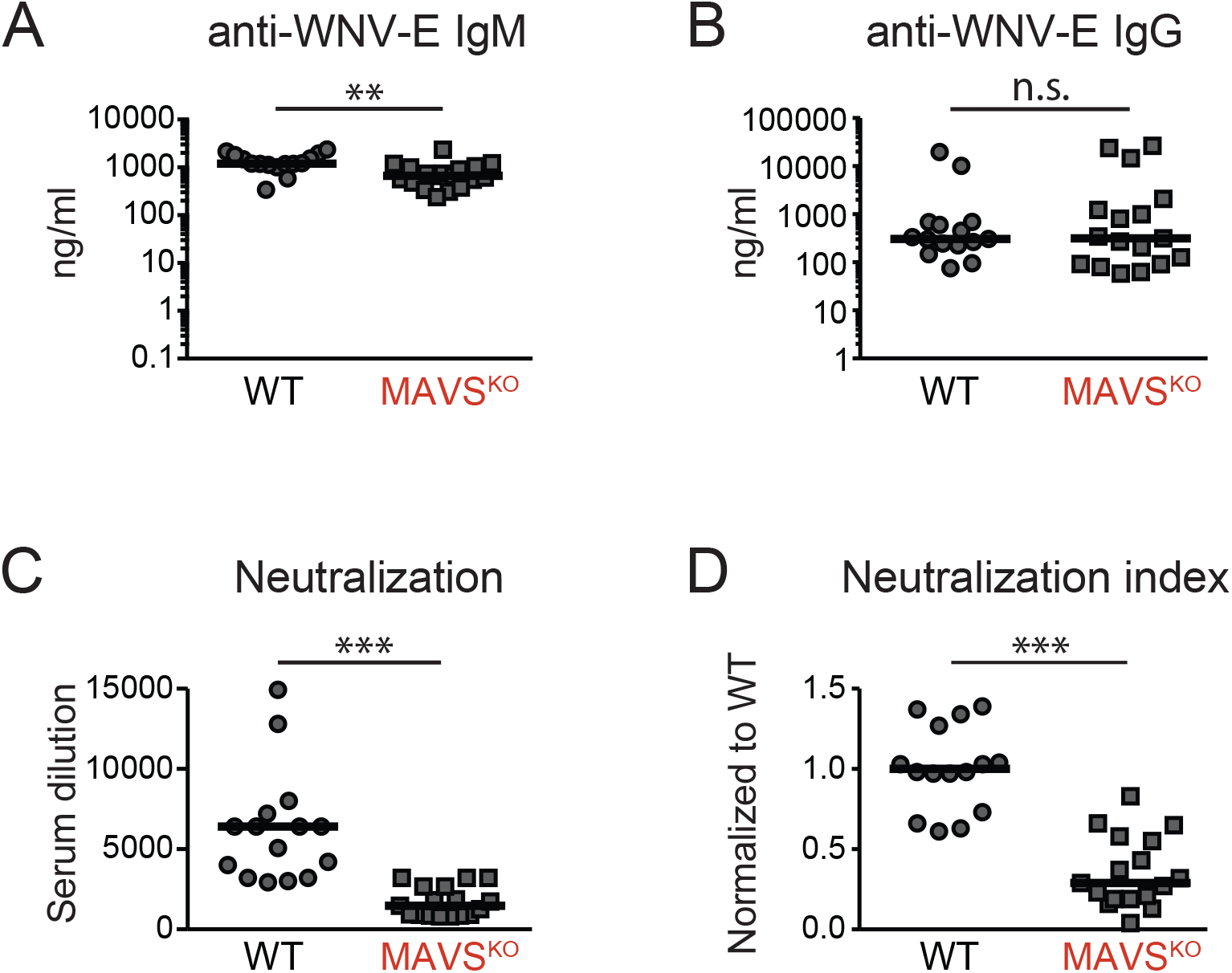
Impaired neutralizing antibody response in RWN-infected MAVS^KO^ mice. **(A, B)** WNV-E-specific IgM (A) and IgG (B) on day 8 after infection with RWN (10^5^ pfu/footpad) as measured by ELISA. **(C)** Virus neutralization by sera of infected MAVS^KO^ and MAVS^WT^ mice. RWN was incubated with serial dilutions of sera prior to infection of Vero cells *in vitro*. The number of infected cells was determined 30 h later by staining with an anti-WNV-E antibody. The reduction of infected cells by 90% was scored (PRNT90). **(D)** Neutralization index on day 8 post infection. The index normalizes virus neutralization to the total amount of WNV-E-specific antibodies in MAVS^KO^ and MAVS^WT^ mice. The index was calculated by dividing the dilution factor (PRNT90) of each mouse by the total amount of WNV-E-specific IgM and IgG of the same mouse. The data were normalized across multiple experiments to MAVS^WT^ mice. Each dot represents one mouse, the lines represent the median. **, p <0.005; ***; p < 0.0005; n.s., not significant; Mann-Whitney test.

Due to the repetitive nature of the envelope proteins on viral surfaces, many viruses can elicit a combination of T-dependent or T-independent antibody responses. Indeed, the primary antibody response to replicating WNV is initially independent of CD4^+^ T cells and becomes mainly T-dependent by day 10. As the kinetics of the antibody response to RWN may differ from that to WNV and MAVS is known to regulate components of the complement cascade, we wanted to ascertain that the observed impairment of virus neutralization by sera from RWN-infected MAVS^KO^ mice on day 8 is due to a defect of the T-dependent antibody response itself. The neutralization defect of sera from RWN-infected MAVS^KO^ mice was complement-independent (Supplementary Fig. 2A-B). Thus, we infected CD4 MHCII^KO^ and CD40^KO^ mice as well as wild-type controls with RWN and measured the ability of the sera from these mice to neutralize the virus. In contrast to sera from wild-type mice, sera from either MHCII^KO^ or CD40^KO^ mice were completely devoid of any neutralizing activity to RWN (Supplementary Fig. 2C-D). Together, our data show that viral neutralization by sera from RWN-infected mice depends on the generation of T-dependent antibody responses and is unlikely to involve significant contributions of innate effector mechanisms such as the production of anti-microbial peptides or altered complement activation. Instead, MAVS regulates the quality of the anti-WNV antibody response.

### RWN infection of MAVS^KO^ mice leads causes an increase in viral RNAs but not antigens

To test the nature of RWN infection in MAVS^KO^ mice compared to MAVS^WT^ mice, we infected mice with RWN and measured first the presence of viral RNA encoding WNV-E in the whole draining lymph nodes (dLNs, here: inguinal and popliteal LNs). We found that the dLNs of RWN-infected MAVS^KO^ mice contained significantly more viral RNA than those from MAVS^WT^ mice (Fig. 2A). As RWN is restricted to a single round of infection, we conclude that MAVS^KO^ mice produce more viral RNA per infected cell without expanding the number of infected cells. RLR-mediated inhibition of protein translation during infection with RNA viruses is thought to occur independently of MAVS and instead may be regulated by the innate signaling adaptor STING (30). We therefore determined whether the elevated levels of viral RNA in MAVS^KO^ mice also translate into more viral proteins in the dLNs. We isolated cells from the dLNs from MAVS^KO^ mice and MAVS^WT^ controls on day 1 post RWN-infection, cultured these cells for 24 hrs *in vitro*, and measured the production of WNV-E during that time period in a flow cytometry-based bead assay. We used samples from naïve mice or assay buffer as negative controls. Cells from both MAVS^KO^ and MAVS^WT^ mice produced significant amounts of WNV-E compared to controls. However, we did not observe significant differences in WNV-E production between MAVS^KO^ and MAVS^WT^ mice, even though we noticed a modest trend towards higher WNV-E production in MAVS^KO^ mice (Fig. 2B-C). Together, our data show that RWN infection of MAVS^KO^ mice leads to the expression of more viral RNA compared to MAVS^WT^ mice but does not significantly alter the levels of viral antigens. We conclude that RWN infection of MAVS^KO^ mice overcomes the caveats associated with replication-competent WNV, namely severe immune-pathology and abundance of viral antigens due to the uncontrolled viral replication in these mice.

**Figure 2.**
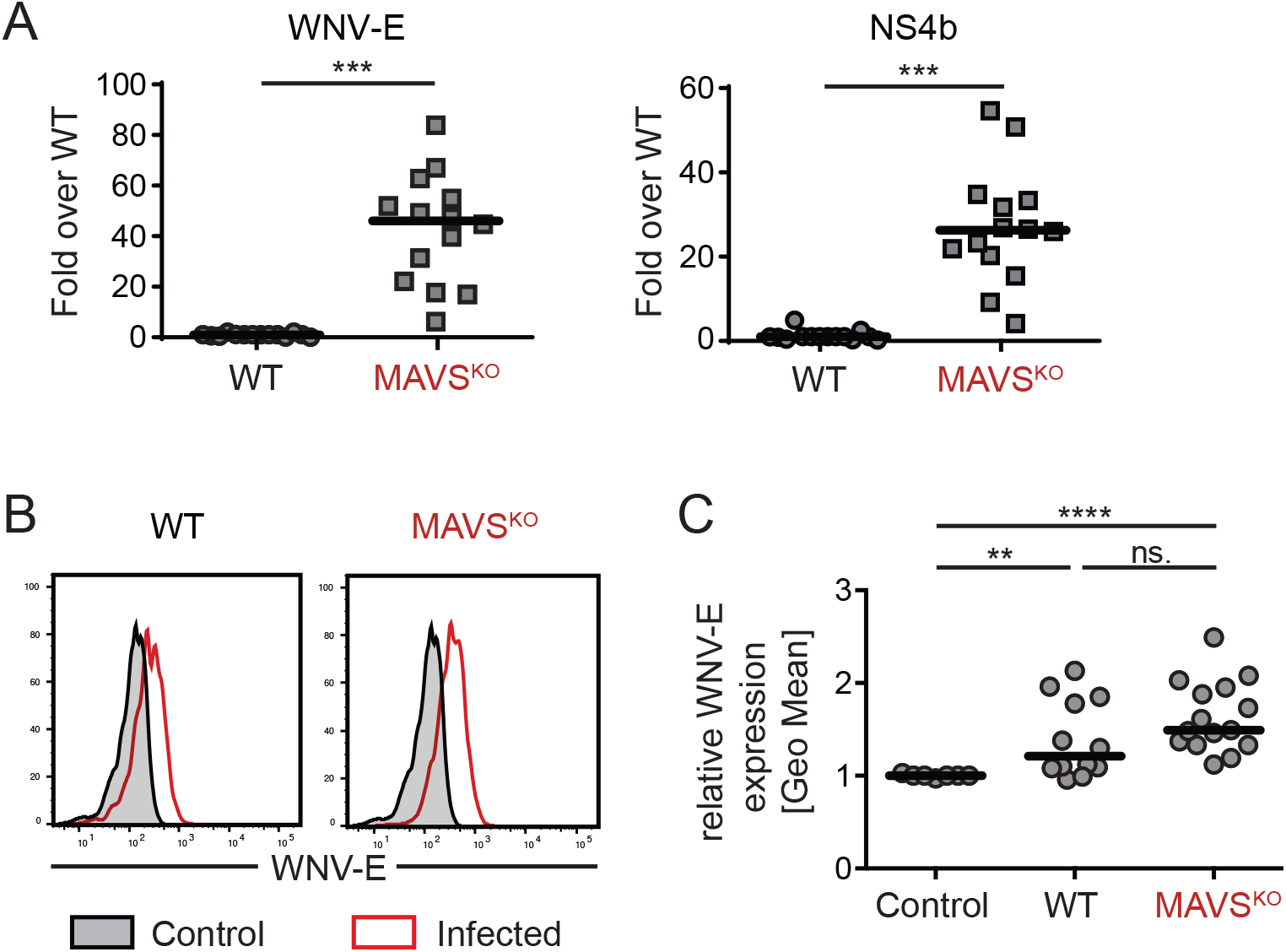
Similar levels of WNV-E protein in RWN-infected MAVS^KO^ and MAVS^WT^ mice. **(A)** Viral RNA levels in the dLNs on day 1 post infection with RWN (10^5^ pfu/footpad) as measured by qPCR using primer pairs located in the WNV-E or NS4b genes of the viral genome. Data were normalized to the RNA level of RWN-infected MAVS^WT^ mice. **(B, C)** Production of WNV-E protein in cells from the dLNs of RWN-infected MAVS^KO^ and MAVS^WT^ mice. Cells from the dLNs were isolated 24 hrs after infection and cultured for an additional 24 hrs *in vitro*. The amount of WNV-E protein in the combined cell lysates and supernatants was quantified by flow cytometry using an anti-WNV-E bead assay. Samples from naïve mice or assay buffer were used as controls. **(B)** A representative experiment is shown. Shaded area represents the background staining of samples from uninfected animals, red lines represent RWN-infected animals. **(C)** Statistical summary of geometric means of multiple independent experiments. The data were normalized to the background staining of uninfected mice in each experiment. Each dot represents one mouse; the line is the median; n.s., not significant, ***, p < 0.0005, Mann-Whitney test.

### Antibody-mediated virus neutralization of RWN-infected mice is independent of antigen loads

It is well-known that high antigen loads can affect the quality of the antibody responses (31, 32). In contrast to infections with replicating WNV, MAVS^KO^ mice infected with RWN do not produce significantly higher amounts WNV-E protein in the dLNs than MAVS^WT^ mice, suggesting that uneven antigen loads are not major drivers of the observed neutralization defect in RWN-infected MAVS^KO^ mice. Nonetheless, as we noticed a trend towards higher levels of WNV-E expression in MAVS^KO^ mice (Fig. 2B, C), we tested whether an increase of the infectious dose of RWN can impact the anti-WNV neutralization index in MAVS^WT^ mice. We first compared MAVS^KO^ mice infected with the standard dose of 10^5^ Pfu per footpad to MAVS^WT^ mice infected with a high dose of 10^6^ Pfu per footpad to ensure that the chosen increased dose for MAVS^WT^ mice leads to similar or higher levels of WNV-E protein in the dLNs. Cells from the dLNs of MAVS^WT^ mice infected with the high dose did indeed express equivalent amounts of WNV-E protein as MAVS^KO^ mice infected with the standard dose (Fig. 3A). Importantly, increasing the infectious dose in MAVS^WT^ mice did not impact neutralization efficiency of the antibody response as this did not negatively impact the neutralization index (Fig. 3B). We thus conclude that the impact of MAVS on the quality of the antibody response in RWN-infected animals is independent of the antigen load.

**Figure 3.**
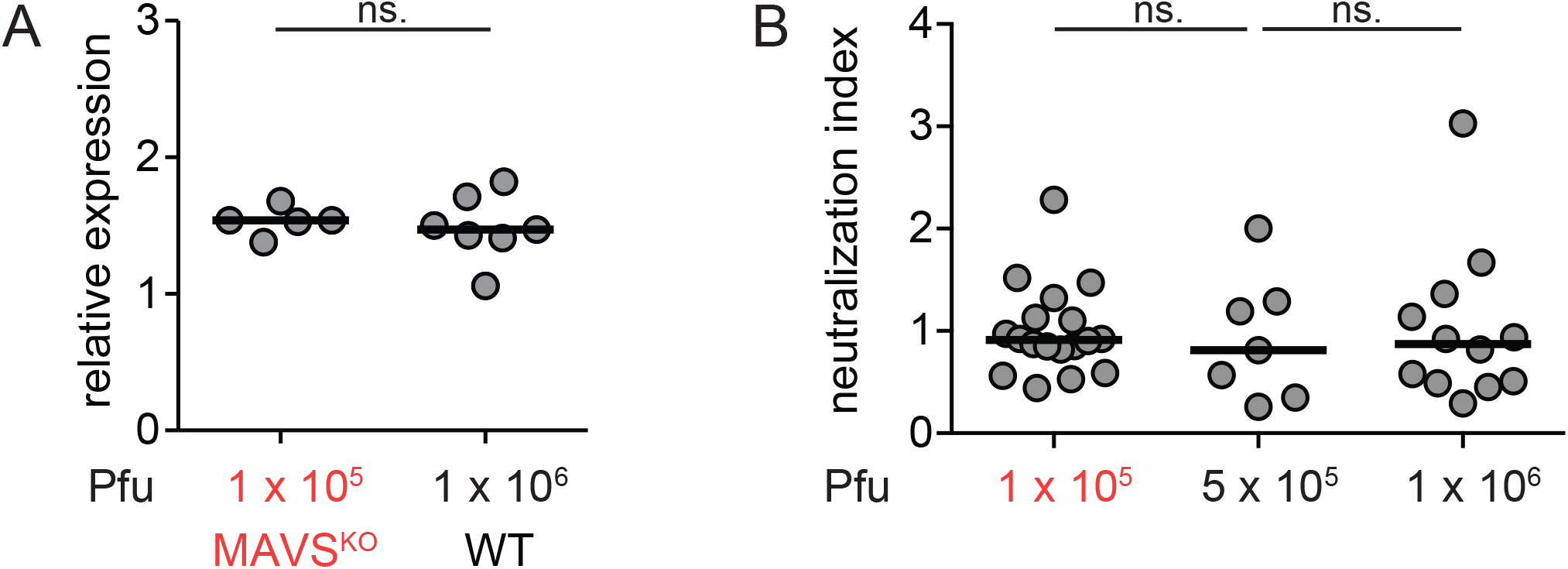
Increased antigen production does not impair virus neutralization. **(A)** Comparison of the viral load in the dLNs of MAVS^KO^ mice infected with 10^5^ Pfu RWN and MAVS^WT^ mice infected with an increased dose of 10^6^ Pfu RWN. Cells from the dLNs were isolated 24 hrs after infection and cultured for an additional 24 hrs *in vitro*. The amount of WNV-E protein in the combined cell lysates and supernatants was quantified by flow cytometry using an anti-WNV-E bead assay. **(B)** Increased doses of RWN infection do not impair virus neutralization in MAVS^WT^ mice. Mice were infected with indicated doses of RWN. The amount of WNV-E-specific IgM and IgG as well as PRNT90 were determined in order to calculate the neutralization index. Shown are the combined data of 3 experiments. Each dot represents one mouse, the line represents the median. ns, not significant; Mann-Whitney test.

### Enhanced Tfh cell and GC B cell response to RWN in MAVS^KO^ mice

In order of gain insights into the potential drivers for the impaired neutralizing antibody response, we next characterized the T and B cell response of RWN-infected MAVS^KO^ mice on the cellular level. MAVS^KO^ mice had normal B cell numbers in the dLNs on day 8 after RWN infection (Fig. 4A). However, germinal center (GC) B cells were more frequent in these mice (Fig. 4B). CD4^+^ T cell numbers were increased in the dLNs of MAVS^KO^ mice (Fig. 4C). CD4^+^ T cells specific for the immuno-dominant WNV epitope E641 were present in similar frequencies in MAVS^KO^ and MAVS^WT^ mice as shown by staining with E641:I-A^b^ MHC class II tetramers (Fig. 4D). However, as the CD4^+^ T cell compartment was enlarged in MAVS^KO^ mice, these mice contained significantly more E641:I-A^b+^ CD4^+^ T cells in the dLNs than MAVS^WT^ controls (Fig. 4D). Similarly, the frequency of CXCR5^+^PD-1^+^ Tfh cells did not change in MAVS^KO^ mice (Fig. 4E) but Tfh numbers were increased (Fig. 4E). Together, our findings may imply that the qualitative defect of the antibody response to RWN in MAVS^KO^ mice is caused by an impaired recruitment of WNV-E-specific B cells with high affinity into the response or their impaired selection into the plasma cell compartment, either because of a defect that acts on B cells directly or a reduced selection pressure due to an increase in Tfh cell number or function (33).

**Figure 4.**
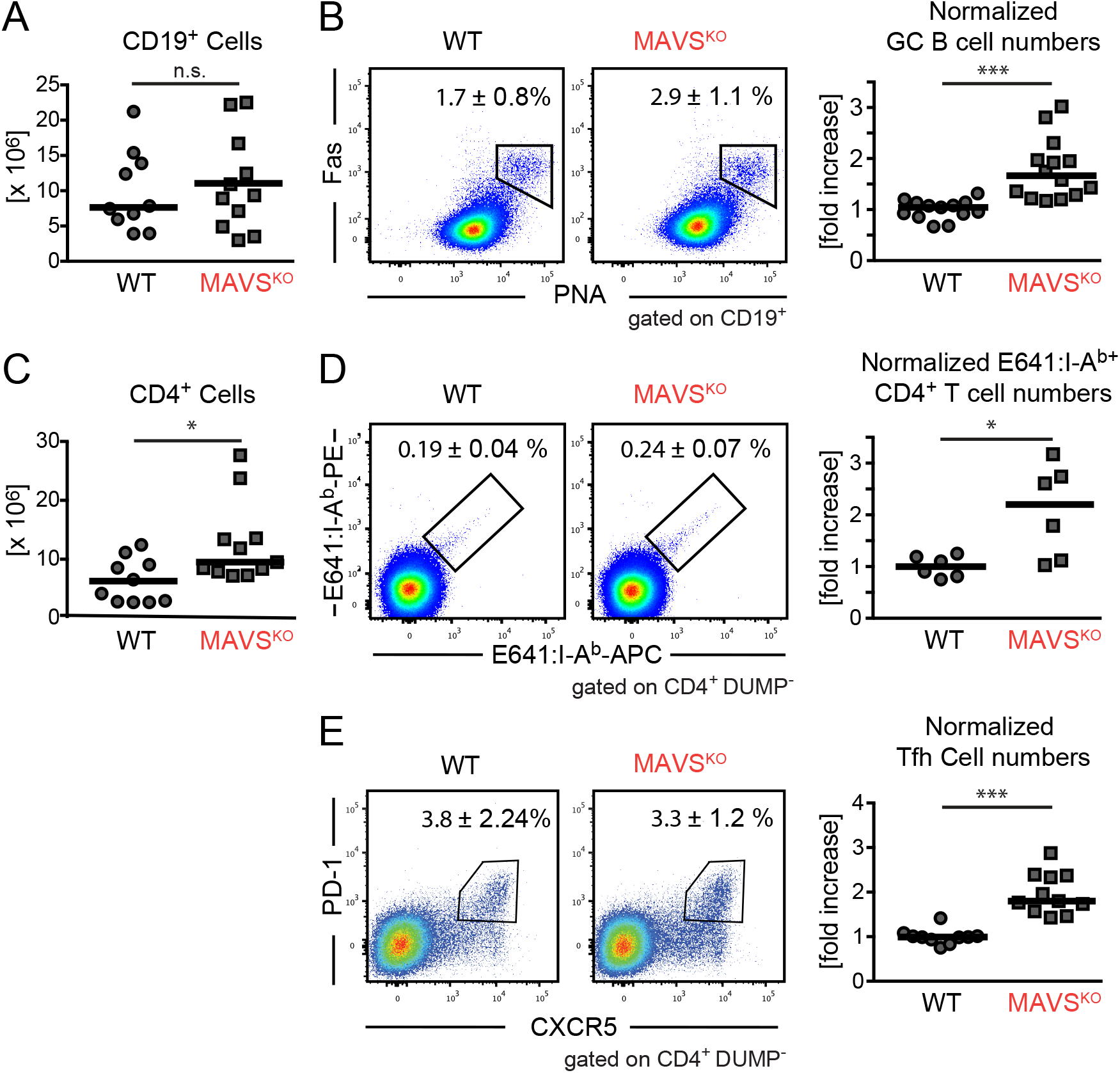
Enhanced GC B and CD4^+^ T cell response to RWN in MAVS^KO^ mice. The cellularity of the dLNs from MAVS^KO^ and MAVS^WT^ mice was analyzed 8 days post RWN infection (10^5^ pfu/footpad) by flow cytometry. **(A)** Total numbers of CD19^+^ B cells. **(B)** Left panels: Frequency of Germinal Center (GC) B cells. Right panel: Total GC B cell numbers of 5 experiments normalized to the average of MAVS^WT^ mice in each experiment. **(C)** Absolute number of CD4^+^ T cells. **(D)** Left panels: Frequency of E641:I-A^b^ class II tetramer^+^ CD4^+^ T cells specific for the immunodominant E641 epitope derived from WNV-E. Right panel: Total E641:I-A^b+^ CD4^+^ T cells numbers of 3 experiments normalized to the average of MAVS^WT^ mice in each experiment. **(E)** Left panels: Frequency of CXCR5^+^ PD-1^+^ T follicular helper (Tfh) cells. Right panel: Total Tfh cell numbers of 5 experiments normalized to the average of MAVS^WT^ mice in each experiment. Frequencies are shown as mean ± SD. Cell numbers: Each dot is one mouse, lines are the medians. *, p < 0.05; ***; p < 0.0005; n.s., not significant; Mann-Whitney test.

### Altered cytokine production in the dLNs of MAVS^KO^ mice

RLRs are major inducers of NF-κB-driven proinflammatory cytokines and type I and type III IFN responses. In fact, the induction of IFNs in particular often depends on the activation of RLRs in infections with RNA viruses. As such changes in the cytokine milieu may alter the generation of T or B cell responses in RWN-infected MAVS^KO^ mice, we determined the expression of a selection of cytokines and IFNs is altered in these mice. We measured the expression of cytokines, IFNs, and interferon-sensitive genes (ISGs) in the whole dLNs of MAVS^KO^ and MAVS^WT^ mice 24 h after RWN-infection by qRT-PCR. We chose this time point because IFN responses peak early in viral infections and innate instruction of CD4^+^ T cells and B cells is thought to occur at that time as well. Unexpectedly, RWN-infected MAVS^KO^ mice expressed more IL-1β, IL-6, and IFN-λin the dLNs than MAVS^WT^ controls, whereas MAVS^KO^ and MAVS^WT^ mice expressed equal amounts of type I IFNs and TNF-α(Fig. 5A-B). Systemic type I IFNs in the serum of RWN-infected mice were only detectable by a type I IFN-sensitive bioassay and were not significantly different between MAVS^KO^ mice and MAVS^WT^ controls, suggesting that IFNs mainly act locally in RWN-infected mice (Supplementary Fig. S3). Finally, ISG expression was unchanged in the dLNs of MAVS^KO^ mice (Fig. 2C). Proinflammatory cytokines remained expressed at the same levels in MAVS^KO^ and MAVS^WT^ mice 48 h post infection, while IFNs and ISGs were, as expected, downregulated (Supplementary Fig. S4). Together, the data suggest that inflammatory mediators are not uniformly dysregulated in MAVS^KO^ mice and that the expression of specific cytokines known for their ability to promote Tfh or B cell immunity such as IL-1β and IL-6 are is increased in these mice (34, 35). Of note, we did not observe that specific cytokines, IFNs, or ISGs were downregulated in MAVS^KO^ mice, suggesting that the expression of these genes is driven by other PRRs than RLRs.

**Figure 5.**
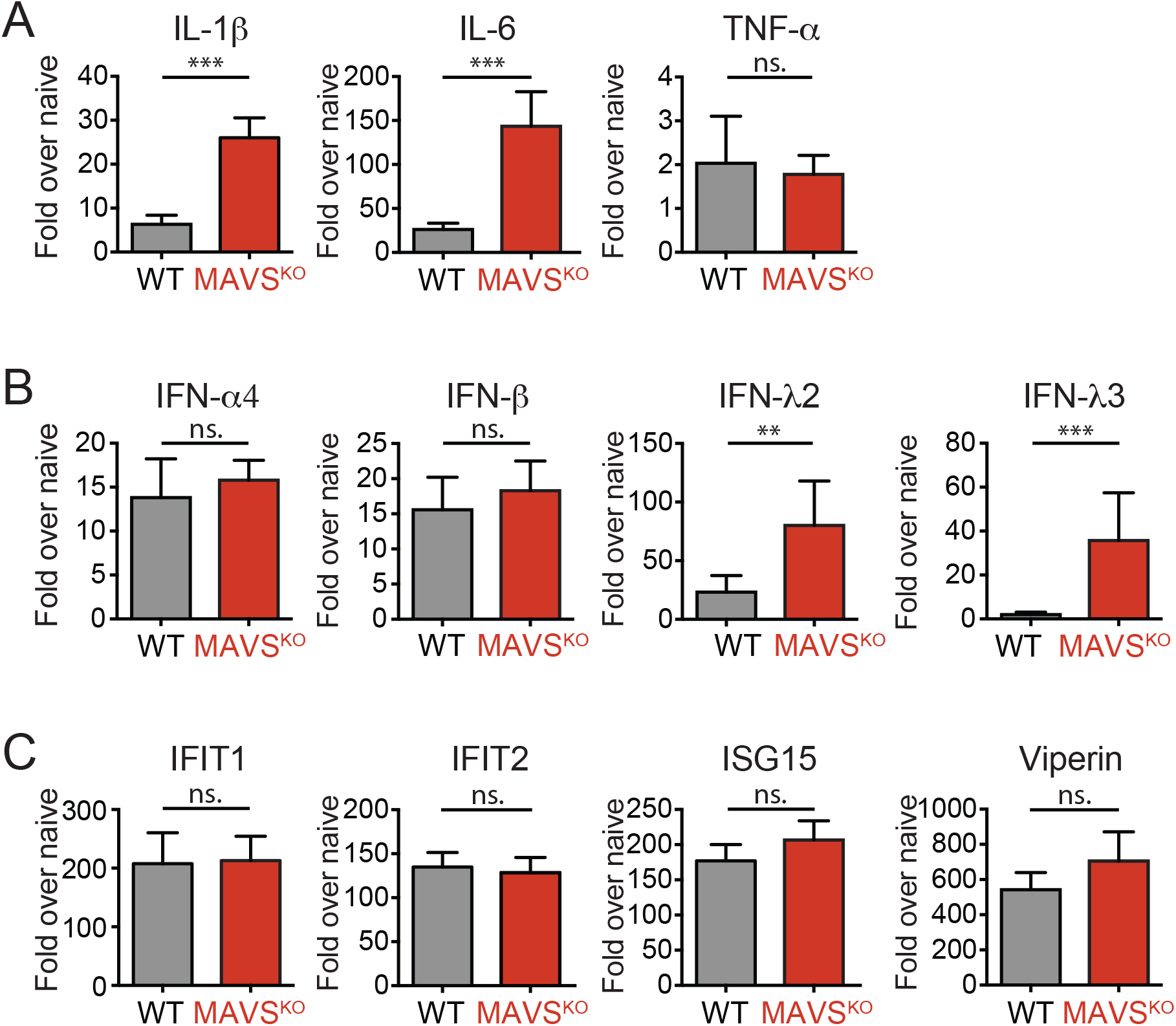
Enhanced production of cytokines and interferon-stimulated genes (ISGs) in the dLNs of MAVS^KO^ mice. **(A)** Expression of IL-1β, IL-6, and TNF-α mRNA RWN-infected MAVS^KO^ and MAVS^WT^ mice. **(B)** Expression of type I and type III IFNs mRNA in RWN-infected MAVS^KO^ and MAVS^WT^ mice. **(C)** Expression of representative ISGs in RWN-infected MAVS^KO^ and MAVS^WT^ mice. **(A-C)** mRNA was isolated from whole dLNs cells of mice 24h after infection with 10^5^ Pfu RWN per footpad and measured by qPCR. Shown is the expression over that of dLNs from naïve WT mice. Expression of GAPDH was used to normalize the samples. Shown are the combined data of 4 experiments using 8-12 mice/genotype **, p < 0.005; ***, p < 0.0005; Mann-Whitney test.

### Antibodies specific for the neutralizing epitope in the WNV-E protein are efficiently generated in MAVS^KO^ mice

Alterations in the Tfh and B cell response of MAVS^KO^ mice (Fig. 4) suggested that the selection of WNV-E-specific B cells is affected in these mice. Although RWN-infected MAVS^KO^ mice produce similar amounts of WNV-E-specific antibodies, they may preferentially generate antibodies that are directed at non-neutralizing epitopes. The major neutralizing epitope of WNV is located in the lateral ridge (LR) epitope of the domain III (DIII) of WNV-E (24, 36, 37). Thus, we used recombinant DIII protein as antigen in ELISAs to test whether the serum of MAVS^KO^ and MAVS^WT^ mice contain similar amounts of DIII-LR-specific antibodies. We also measured the antibody titers specific for a mutant form of DIII with an altered LR epitope (DIII-K307E/T330I) that abrogates the binding of DIII-LR-specific antibodies. We found that the levels of both IgM and IgG specific for anti-DIII remained unchanged MAVS^KO^ mice (Fig. 6A). The same was true for anti-DIII-K307E/T330I antibodies in these mice (Fig. 6B). Consistent with previous findings, the anti-DIII-K307E/T330I antibody titers were much lower than those specific for wild-type DIII, suggesting that the majority of anti-DIII antibodies in both MAVS^KO^ and MAVS^WT^ mice are directed against the neutralizing DIII-LR epitope (25). Importantly, the ratio of antibodies bound to DIII versus DIII-K307E/T330I was also similar in MAVS^KO^ and MAVS^WT^ mice (Fig. 6A, B). A recalculation of the neutralization index (Fig. 1D) with the anti-DIII antibody titers reinforced the notion that the neutralizing Ab response is compromised in MAVS^KO^ mice (Supplementary Fig. S5). Together, these findings demonstrate that MAVS^KO^ mice produce antibodies against WNV with similar specificities as MAVS^WT^ mice and a lack of antibodies against the neutralizing DIII-LR epitope is not responsible for the neutralizing defect of MAVS^KO^ mice.

**Figure 6.**
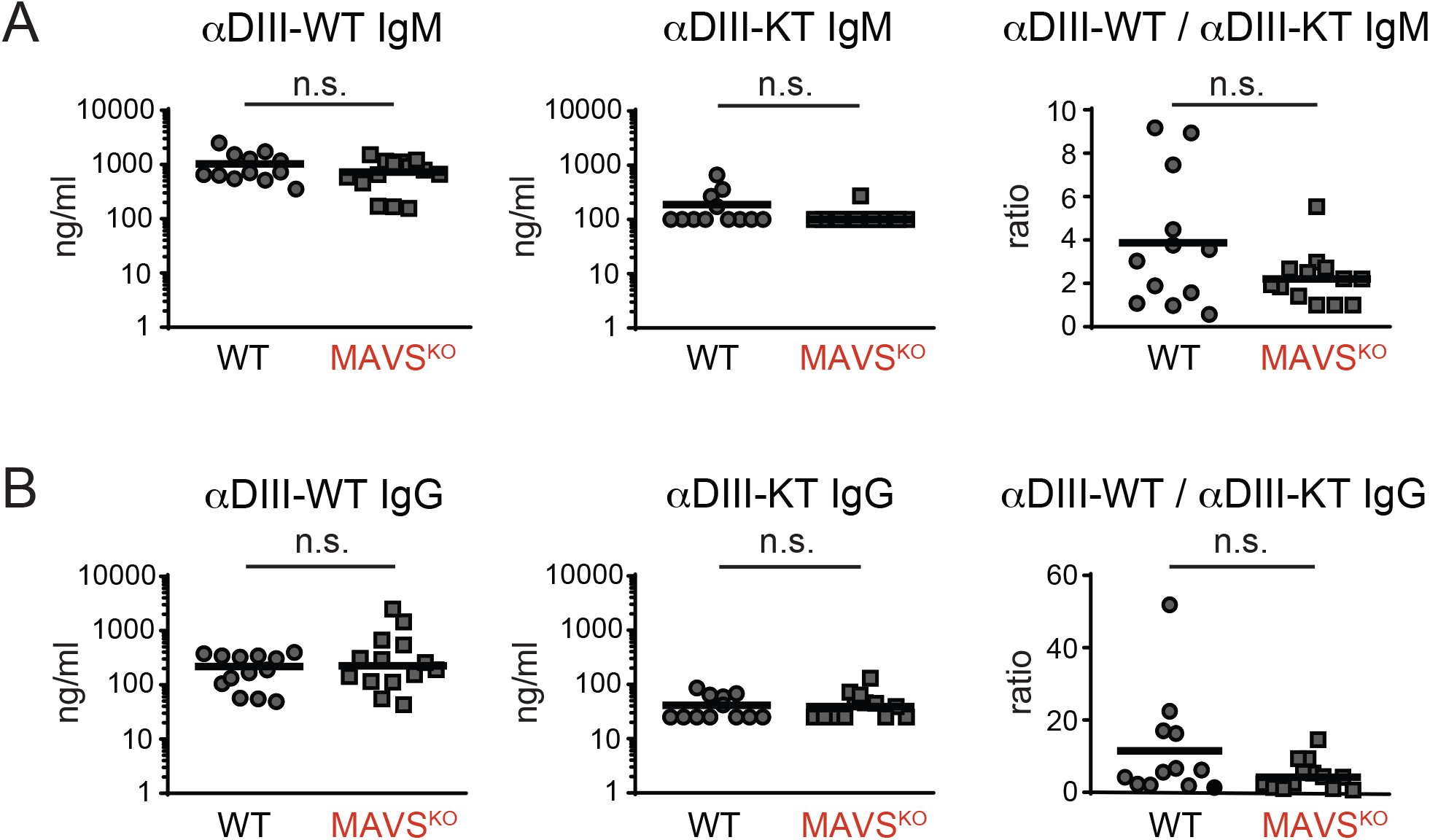
Normal specificity of the antibody response of MAVS^KO^ mice to the neutralizing lateral ridge (LR) epitope in the WNV-E DIII domain. **(A, B)** The antibody response of RWN-infected MAVS^KO^ and MAVS^WT^ mice was measured 8 days post infection. Shown are the titers of IgM **(A)** and IgG **(B)** specific for the WT DIII domain or the mutant form DIII-KT containing loss-of-function mutations in the neutralizing LR epitope (DIII-K307E/T330I). Ratios represent the excess of antibodies directed at the neutralizing epitope over the amount of non-neutralizing antibodies directed at the DIII domain. Combined data of 4 experiments. Each data point is one mouse, the line is the median; n.s., not significant; Mann-Whitney test.

### Decreased affinity of anti-DIII antibodies in MAVS^KO^ mice

Given the unchanged specificity of the antibody response in MAVS^KO^ mice, we hypothesized that the defect in virus neutralization of MAVS^KO^ is caused by a lower avidity of the neutralizing antibodies in these mice. To test this, we measured the avidity of the antibodies against the DIII domain by ELISA in the presence of increasing amounts of NaSCN to enhance the stringency of antibody binding. For IgM, we also reduced the binding avidity by diluting the recombinant DIII antigen with BSA. Consistent with previous results (Fig. 1), we did not find any differences in the total anti-DIII IgM titers between MAVS^KO^ and MAVS^WT^ mice (Fig. 7A). However, upon dilution of DIII with BSA and increasing concentration of NaSCN, we observed a successive reduction of the anti-DIII IgM titers from sera of RWN-infected MAVS^KO^ mice while sera from MAVS^WT^ mice retained their ability to bind to DIII significantly better (Fig. 7A). The quantification of these results, in which we expressed the amount of high avidity anti-DIII IgM as percentage of total anti-DIII in each mouse, confirmed this result (Fig. 7B). In contrast to the anti-IgM response of MAVS^KO^ mice, we did not observe any differences in the anti-DIII IgG response of these mice compared to MAVS^WT^ controls (Fig. 7C-D). The latter observation was consistent with the notion that the primary antibody response to WNV is dominated by IgM whereas somatically mutated high-affinity IgG responses emerge late in the primary response and are mainly required for protection from secondary challenges (38–40). Together, these results show that MAVS-deficiency results in a qualitatively inferior primary antibody response due to a reduced affinity to the neutralizing LR epitope of WNV-E protein. In light of the defective IgM response, which consists mostly of unmutated antibodies, these findings also imply that MAVS directly affects the recruitment of B cells into the immune response.

**Figure 7.**
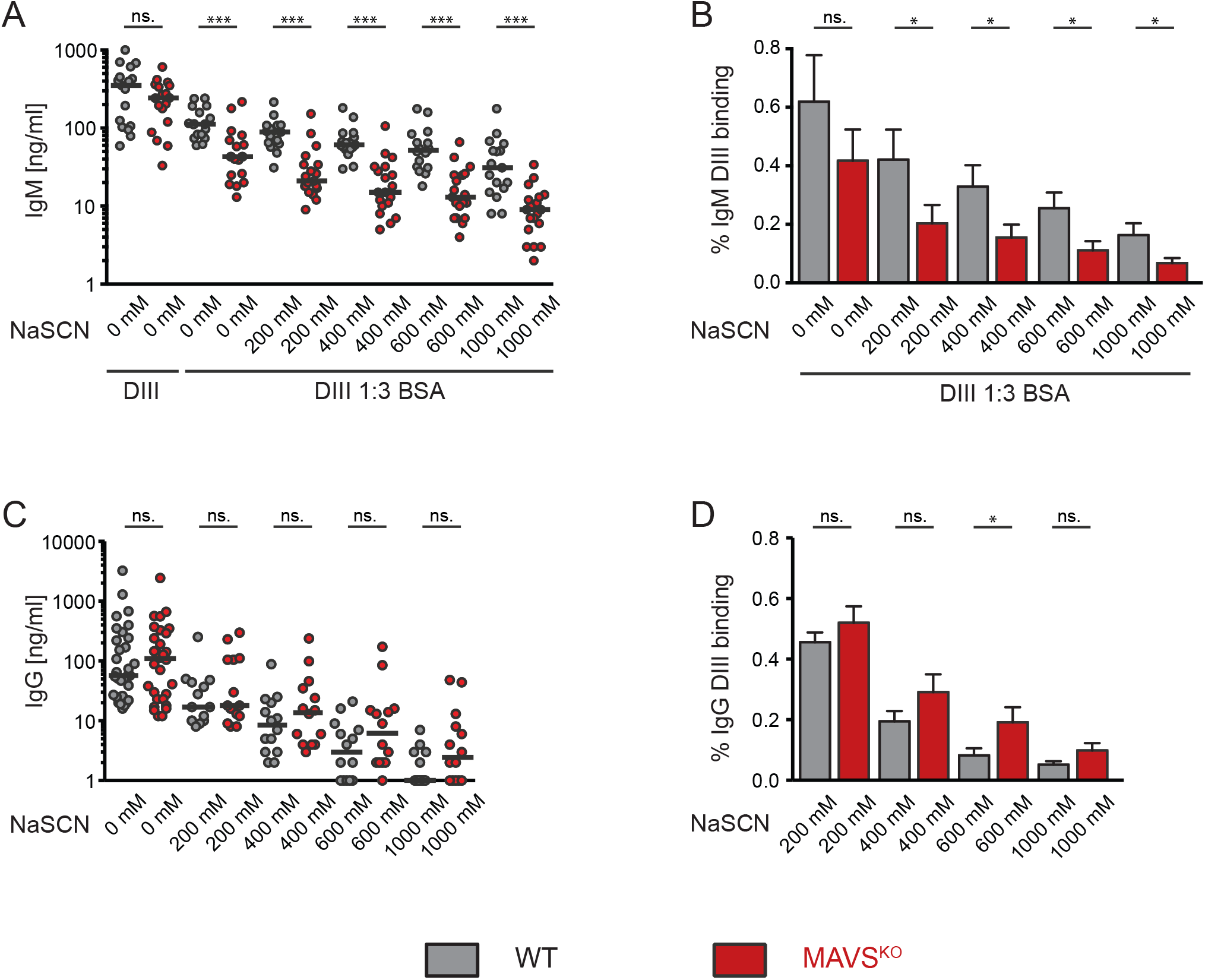
Impaired affinity of the IgM response to the DIII domain of WNV-E protein in RWN-infected MAVS^KO^ mice. **(A, B)** DIII-specific IgM titers from RWN-infected mice were measured by ELISA in the presence of BSA to reduce binding avidity and increasing amounts of NaSCN to enhance stringency of binding. Recombinant DIII protein was used as antigen. Shown are the absolute titers (A) and the titers as fraction of the total amount of DIII-specific antibodies in the absence of BSA and NaSCN for each mouse (B). **(C, D)** Same as before for DIII-specific IgG. Shown are the combined data of 4 experiments. Each dot represents one mouse, the line is the median. *, p < 0.05; ***, p < 0.0005; n.s., not significant; Mann-Whitney test.

## Discussion

In contrast to the better-known functions of transmembrane PRRs, the mechanisms by which cytosolic PRRs regulate adaptive immunity are still poorly understood. Previous work suggested that the detection of viral RNA by RLRs and their signaling adaptor MAVS is an important step in the control of adaptive immunity to pathogenic WNV (17, 18, 41). While these results seemingly established a link between MAVS signaling and the generation of protective adaptive immune response, it was difficult to attribute a specific role to MAVS in the regulation of such responses as the unrestricted viral replication in the absence of MAVS-dependent innate immune defenses results in strongly elevated levels of viral antigens and RNAs as well as severe pathology and death (17). To overcome these complications, we have used a replication-incompetent mutant of a pathogenic WNV called RepliVAX-WN (RWN) to study the functions of MAVS in anti-viral humoral immunity.

Although the RLR/MAVS signaling pathway is known to regulate the release of serum components such as specific members of the complement cascade, we show here that the MAVS-dependent effects on WNV neutralization depend on the presence of T-dependent antibodies, consistent with the previously observed important contribution of T-dependent antibodies to the overall humoral immune response to WNV (42, 43). MAVS-deficiency did not alter the overall nature of the antibody response as both the choice of immunoglobulin isotypes and, importantly, the choice of WNV-E epitopes remained the same. Instead, the lack of MAVS reduced the overall affinity of the IgM response, the main isotype responsible for protection during primary infection, to the neutralizing LR epitope of the DIII domain despite otherwise normal anti-DIII antibody titers (36, 37, 44–48). This feature is the likely cause for the impaired WNV neutralization in MAVS^KO^ mice, as antibody affinity defines the antibody occupancy rate on the WNV virion and thus is a major factor that determines WNV neutralization (49).

High antigen loads and the associated abundant formation of immune complexes have been recognized since the early days of B cell immunology for their potential to impair the overall affinity of the antibody response (31, 32). While such a scenario may contribute to the impaired antibody response in the context of uncontrolled replication of pathogenic WNV, it is an unlikely factor in infections of MAVS^KO^ mice with RWN. Here, the replication-incompetent nature of this WNV mutant results in a similar cellular tropism in mice as WT controls, which differs from WNV-infected mice with deficiency in type I IFN responses (50). Despite elevated levels of viral transcripts in the dLNs, we observed only a modest increase of WNV-E protein production in MAVS^KO^ mice early in the infection. These findings are consistent with a recent study indicating that the RIG-I/MDA5-mediated suppression of protein translation following infection with RNA viruses depends on the signaling adaptor STING instead of MAVS (30). The mere increase in the infectious dose did not alter the quality of the neutralizing antibody response. Thus, the defect of the neutralizing antibody response in RWN-infected MAVS^KO^ mice is not a consequence of a fundamental change in the antigen load and instead is likely caused by an altered immune regulation by cytokines or other signaling molecules in the absence of MAVS. Consistent with this view, a recent study of MAVS^KO^ mice infected with a replicating non-pathogenic strain of WNV did not impair the quality of the antibody response, even though these mice exhibited increased viral titers (51).

At the present time, it remains unclear how MAVS regulates the quality of the antibody response. Although we did not resolve the cytokine profile of the individual cell populations in the dLNs, our data nonetheless indicate that several cytokines are upregulated in RWN-infected MAVS^KO^ mice, presumably through the activity of other PRRs such as TLRs. One possibility may be that the altered cytokine environment in the dLNs of RWN-infected MAVS^KO^ mice directly influences the ability of B cells to become activated. Indeed, both type I and type III IFNs can promote B cell responses directly by facilitating B cell receptor activation or indirectly by regulating the B cell-intrinsic activity of TLR7 and other PRRs (52–54). Such signals may facilitate the activation of low affinity B cell that usually would be prevented from participating in the response. In this context, it is interesting that MAVS affects the quality of the neutralizing antibody response to pathogenic strains of WNV, whereas it does not seem to play the same function in infections with non-pathogenic strains (51). A key difference between these viral isolates is their divergent interference with the IFN signaling pathway, further pointing towards a function of IFNs in the regulation of the anti-WNV antibody response (55).

An additional possibility is that altered cytokine profiles may lead to the promotion of an enhanced Tfh cell response. Indeed, we observed increased numbers of antigen-specific CD4^+^ T cells as well as Tfh cells in RWN-infected MAVS^KO^ mice. Here, the observed upregulation of IL-1 and IL-6 may help CD4^+^ T cells to overcome Treg-mediated suppression and promote their differentiation into more effective Tfh cells (35, 56, 57). The consequence of this scenario may therefore be a reduced competition of cognate B cells for Tfh cell help that leads to the entry of low affinity B cells into the immune response. Such a checkpoint is usually recognized in the context of a GC response. Here, the selection of high affinity B cells depends on their more successful access to limited Tfh cell help and becomes less stringent when T cell help is abundant (33, 58). However, a similar checkpoint is thought to occur already at the T-B border before cognate B cells re-enter the B cell follicle (59). The reduced affinity of neutralizing IgM in MAVS^KO^ mice could therefore support the idea that MAVS already prevents the recruitment of low affinity B cells into the anti-WNV immune response at this early stage, thus facilitating the production of an antibody response with overall higher affinity to the neutralizing DIII-LR epitope. IgM is the major isotype responsible for humoral immunity during primary WNV infection, whereas class-switched and somatically mutated IgG antibodies are thought to be more relevant for secondary WNV infection (39). RLR ligands can promote increased immunogenicity and affinity maturation of IgG responses in protein immunizations (60). Although we noted enlarged Tfh cell and GC B cell compartments in MAVS^KO^ mice, we did not observe a reduction of the affinity of DIII-LR-specific IgG antibodies in the mice titers in contrast to IgM. Such a lack of phenotype in the IgG response may not be surprising because RWN-infected cells and MAVS-dependent signals disappear quickly from the draining lymph after infection in our experimental system. Nonetheless, our findings provide a mandate to explore the regulation of GC responses by MAVS in more detail using replicating pathogenic WNV strains.

Regardless of the specific circumstances that promote the production of low affinity antibodies in RWN-infected MAVS^KO^ mice, it is likely that this effect is caused by an absence of MAVS in myeloid cells. This argument rests on the finding that myeloid cells are much more frequently infected by WNV (50). This view is also attractive because myeloid cells and DCs in particular orchestrate the early events of CD4^+^ T cell responses. Recent results with extracellular bacterial infections demonstrate that phagocytosed microbial RNA can be sensed by a combination of transmembrane PRRs such as TLR3 and cytosolic PRRs such as NLRP3 that cooperate to induce IL-1 and type I IFN for the regulation of Tfh responses (8–10). Such regulatory circuits have been shown to affect primarily the magnitude of the antibody response, whereas our study demonstrates a role for the sensing of microbial RNA in the quality of the antibody response. Nonetheless, these findings may therefore set a precedent for the general recognition of microbial RNAs by phagocytes and provide a conceptual basis for the understanding of RLR signaling pathway in the regulation of antibody responses. Future experiments will address the questions about the origin and nature of the MAVS-dependent signals required for the regulation of the antibody response to WNV.

## Acknowledgments

We thank Drs. Deepta Bhattacharya, Michael Kuhns, and Janko Nikolich-Žugich for critical reagents, technical help, and overall comments and suggestions. This work was supported by the Arizona Biomedical Research Foundation and the National Institutes of Health through R56AI130044.

## Supplementary Figures

**Supplementary Figure S1.**
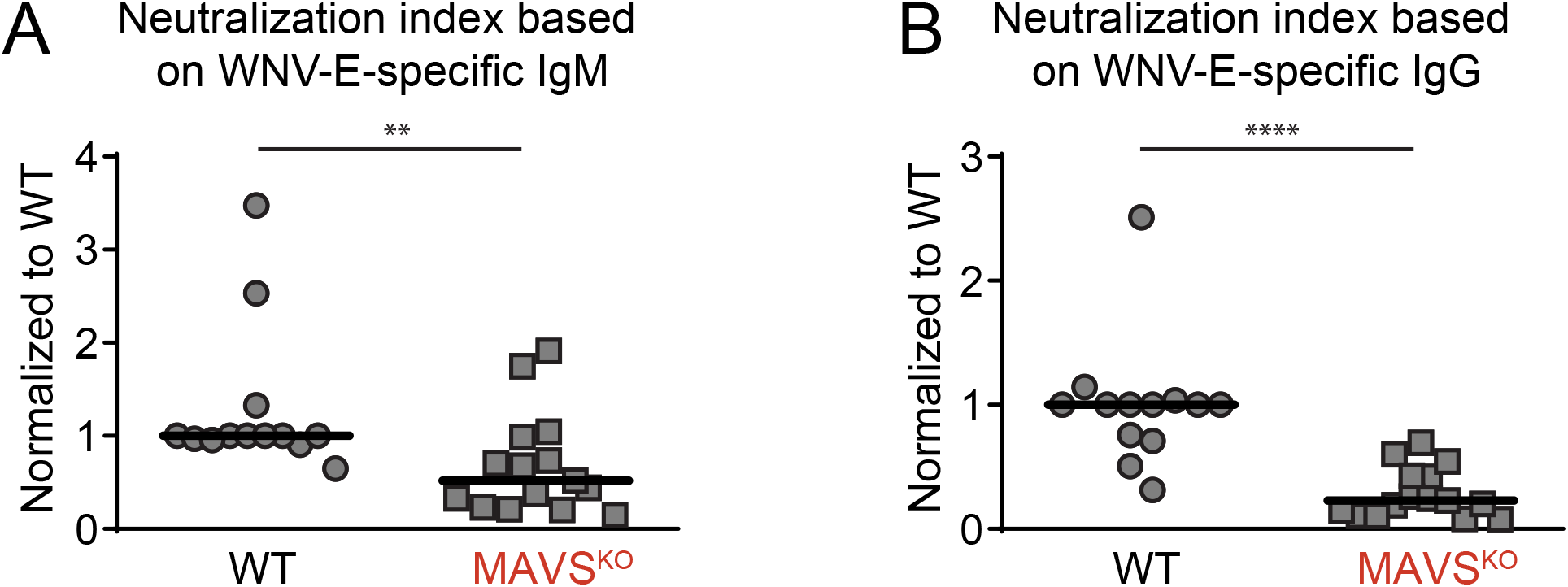
Defective neutralization by serum from RWN-infected MAVS^KO^ mice. **(A-B)** Neutralization index calculated based on serum-specific levels of WNV-ENV-specific IgM **(A)** or IgG **(B)** titers on day 8 after infection. The data were normalized across multiple experiments to WT mice. Each dot represents one mouse, the lines represent the median. **, p <0.005; ****; p < 0.00005; n.s., not significant; Mann-Whitney test.

**Supplementary Figure S2.**
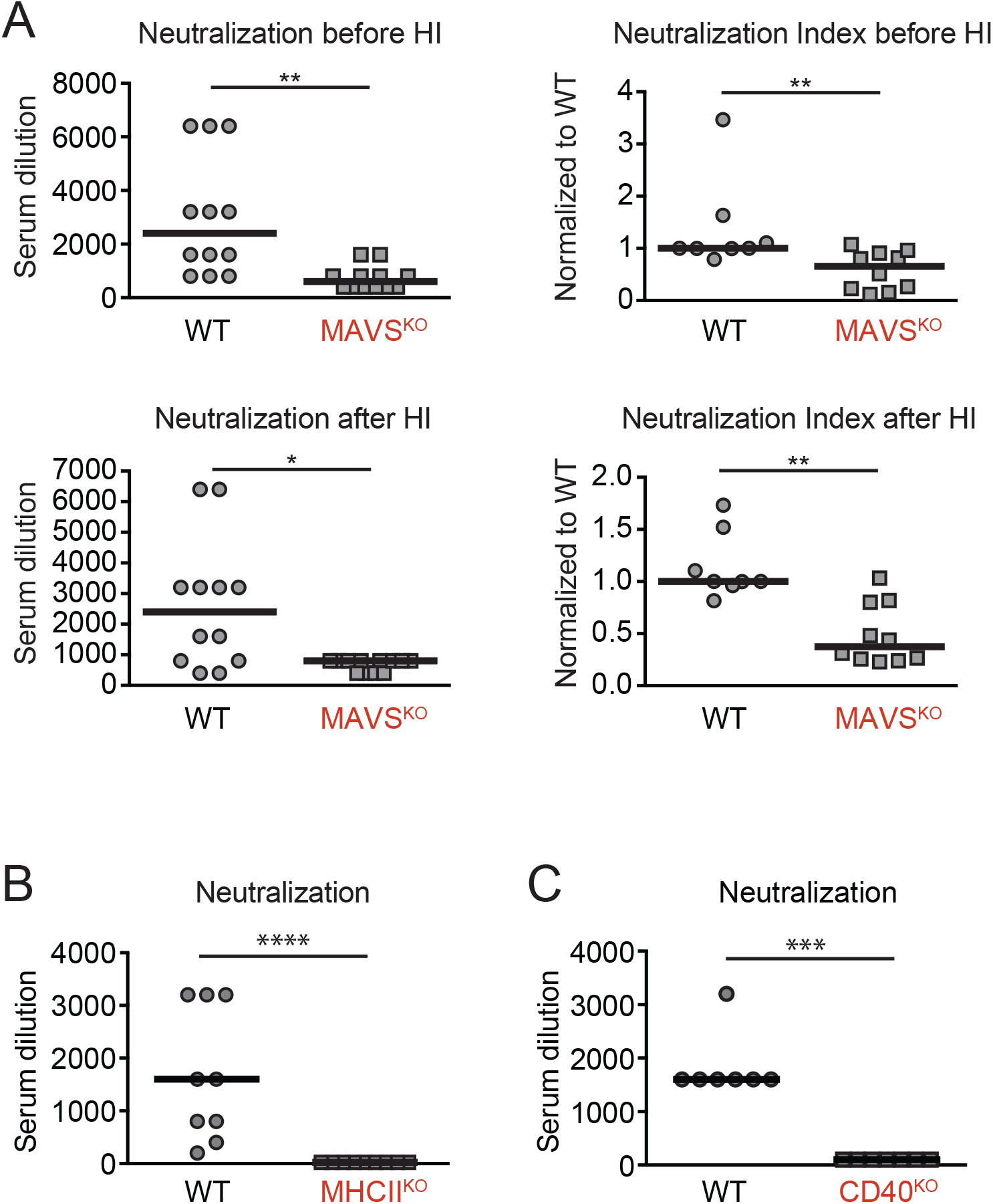
Serum neutralization of RWN is mediated by a T-dependent antibody response. **(A)** Virus neutralization before and after heat-inactivation (HI) to exclude complement-mediated effects. Shown are the dilution factors that resulted in a 90% reduction of infection of target cells with RWN *in vitro* (PRNT90) and the neutralization index that accounts for the anti-WNV-E IgM and IgG titers in each mouse. **(B)** MHCII^KO^ and **(C)** CD40^KO^ mice as well as WT controls were infected with 10^5^ Pfu of RWN in the footpads. Serum was collected 8 days later to measure the dilution factor Shown are the combined data of two independent experiments. Each dot is one mouse. *, p < 0.05; **, p < 0.005; ***, p < 0.0005; ****, p < 0.00005; Mann-Whitney test.

**Supplementary Figure S3.**
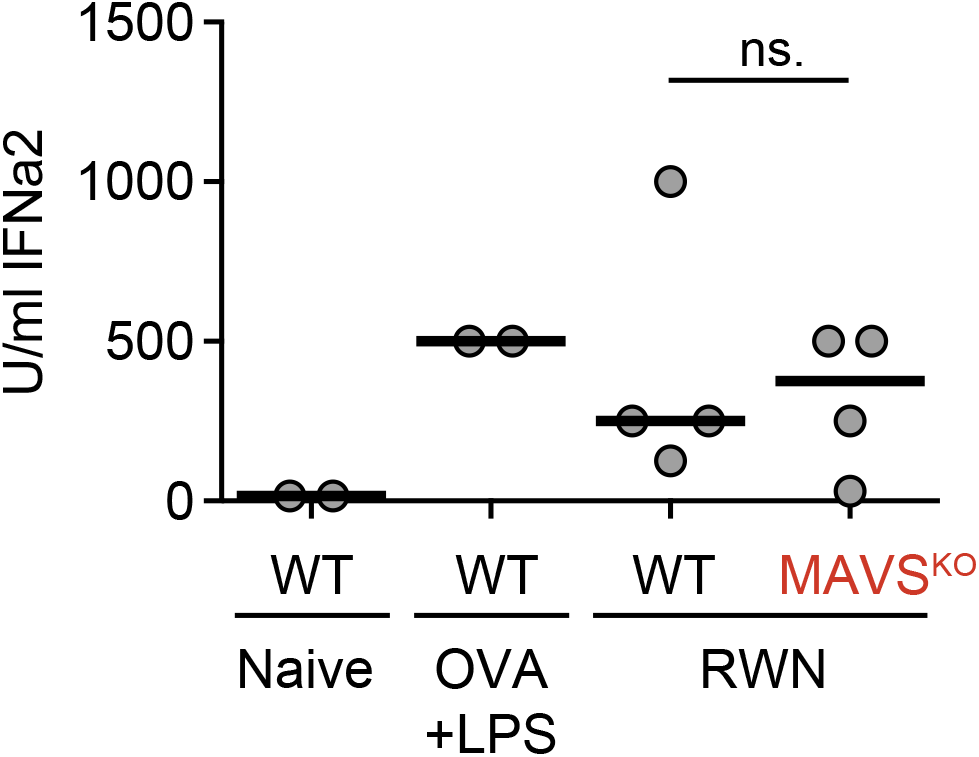
Normal production of systemic type I IFN in the serum of RWN-infected MAVS^KO^ mice. Serially diluted serum samples from day 1 of RWN-infected mice were used to protect L929 cells from the cytopathic effects (CPE) of vesicular stomatitis virus (VSV) *in vitro*. Samples from mice immunized with OVA + LPS were used as positive controls. All samples were compared to samples treated with increasing doses of recombinant IFN-α2 as standards. Shown are the combined data from two experiments. Each dot represents one mouse. n. s., not significant; Mann-Whitney test.

**Supplementary Figure S4.**
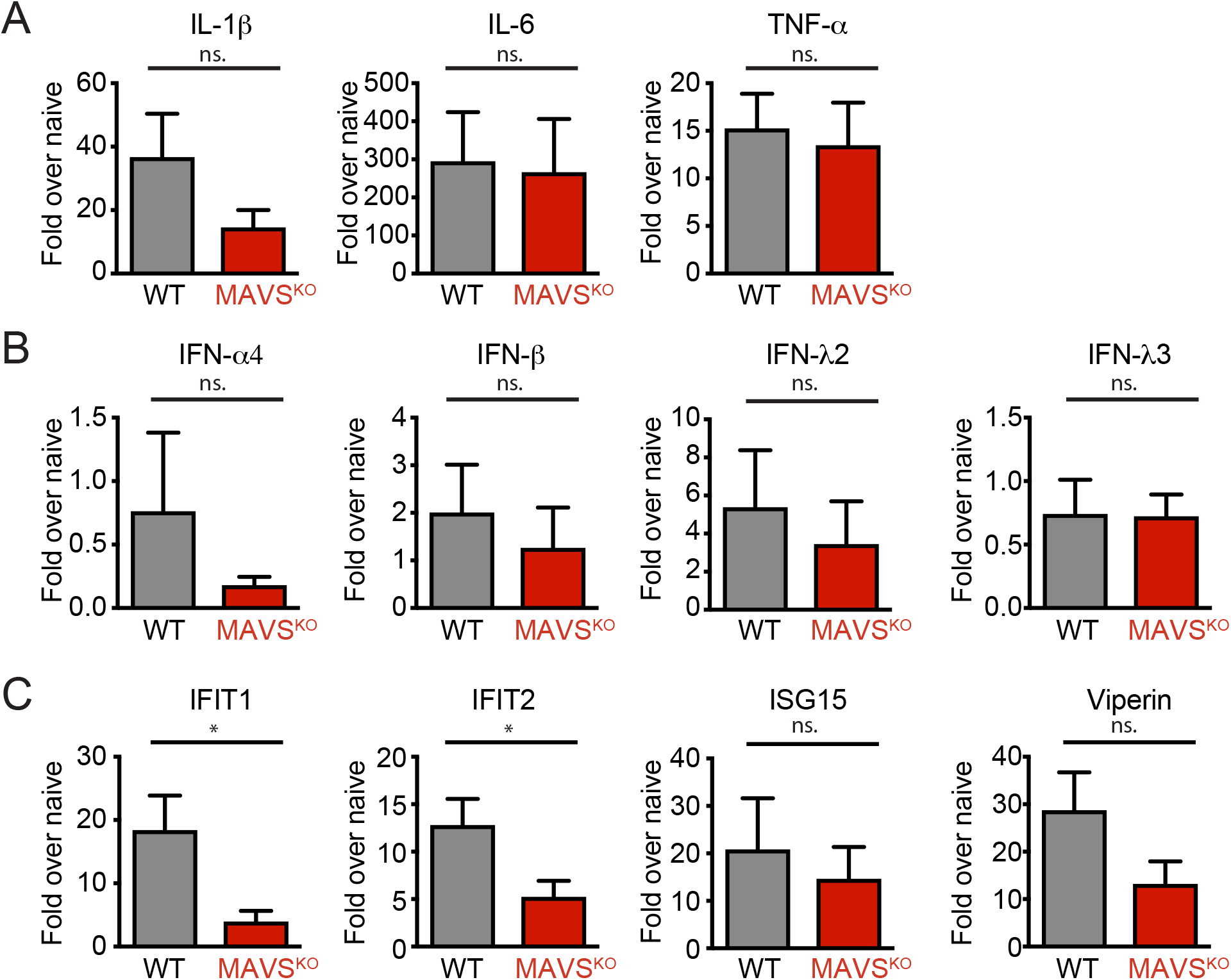
Production of cytokines and interferon-stimulated genes (ISGs) in the dLNs of MAVS^KO^ mice on day 2 post RWN infection. **(A)** Expression of IL-1β, IL-6, and TNF-α mRNA RWN-infected MAVS^KO^ and WT mice. **(B)** Expression of type I and type III IFNs mRNA in RWN-infected MAVS^KO^ and WT mice. **(C)** Expression of representative ISGs in RWN-infected MAVS^KO^ and WT mice. **(A-C)** mRNA was isolated from whole dLN cells of mice 24h after infection with 10^5^ Pfu RWN/footpad and measured by qPCR. Shown is the expression over that of dLNs from naïve WT mice. Expression of GAPDH was used to normalize the samples. Shown are the combined data of 4 experiments using 8-12 mice/genotype **, p <0.005; ***, p < 0.0005; M-W test.

**Supplementary Figure 5.**
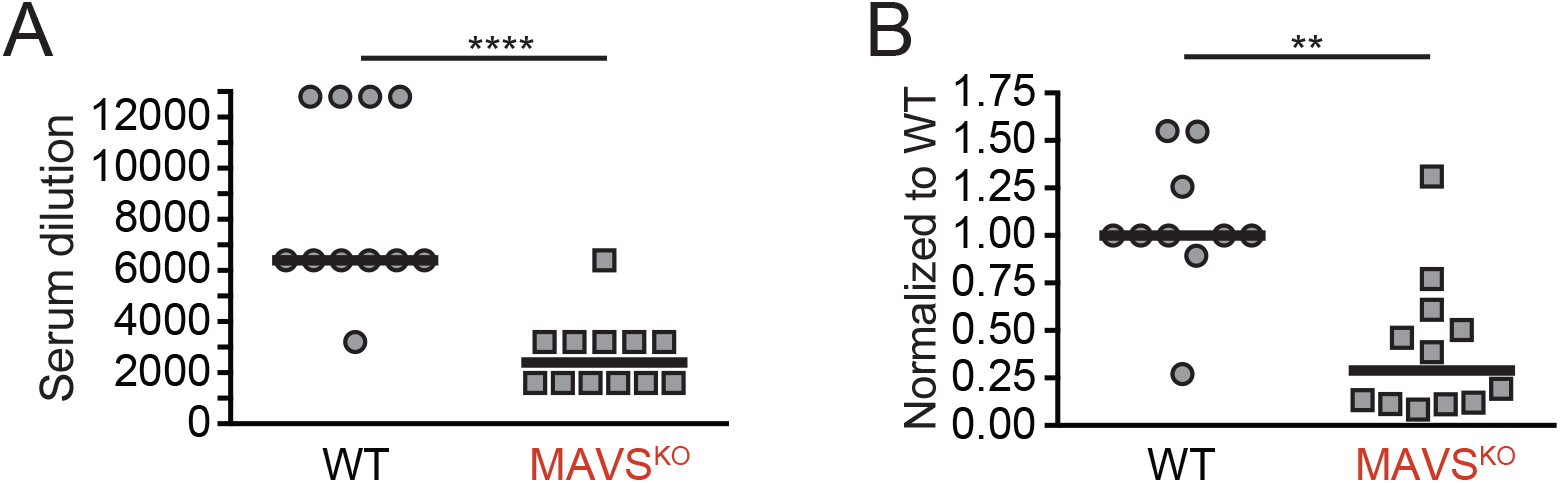
Neutralization index for sera of RWN-infected MAVS^KO^ mice based on the titers of DIII-specific antibodies. **(A)** Virus neutralization with sera from MAVS^KO^ and MAVS^KO^ mice. (B) Neutralization index for the same mice based on the DIII-specific IgM and IgG titers (Dilution factor divided by the total amount of DIII-specific IgM and IgG). Each dot represents one mouse, the line is the median. Shown are the combined data of two experiments. **, p <0.005; ****, p <0.00005; Mann-Whitney test.

